# Chemotherapy induced immunogenic cell death alters response to exogenous activation of STING pathway and PD-L1 immune checkpoint blockade in a syngeneic murine model of ovarian cancer

**DOI:** 10.1101/824094

**Authors:** Sarah Nersesian, Noor Shakfa, Nichole Peterson, Thiago Vidotto, Afrakoma AfriyieAsante, Elizabeth Lightbody, Madhuri Koti

## Abstract

Poor response to platinum/taxane-based chemotherapy has remained a major hurdle in the management of high grade serous carcinoma of the ovary (HGSC). Recurrent HGSC is often treated with liposomal doxorubicin as a second line chemotherapy. Unfortunately, HGSC patients have not benefited from immunotherapies targeting the PD-1/PD-L1 immune checkpoint axis. In a pre-clinical study evaluating the efficacy of a “Stimulator of Interferon Genes” (STING) agonist, we demonstrated the synergistic potential of STING pathway activation in enhancing response to carboplatin chemotherapy and sensitization to PD-1 immune checkpoint blockade (ICB). Since carboplatin and doxorubicin exhibit distinct immunogenic cell death (ICD) inducing potential, we investigated the chemotherapy specific effect in the magnitude of response to exogenous STING pathway activation. Immunocompetent C57/BL6 mice were implanted with ID8-*Trp53^−/−^* cells followed by treatment with carboplatin or doxorubicin. Towards rationalized addition of STING agonist with or without PD-L1 blockade, we first determined the expression of 60 known ICD associated genes at an early time point following the initial treatment with carboplatin or doxorubicin with or without STING agonist. Doxorubicin treated tumours showed significantly higher expression of ICD genes, *Cxcl10, Cd274, Isg15, Psmb9* and *Calr*. Expression changes were further amplified following the addition of STING agonist. Significantly higher expression of *Cxcl10* and *Isg15* were observed in the doxorubicin + STING agonist treated mice compared to carboplatin + STING agonist combination. Interestingly, *Ccl5* gene expression was higher in the tumours from carboplatin or carboplatin and STING agonist combination treated mice compared to those treated with doxorubicin. Plasma cytokine analysis showed distinct profiles of CXCL10, CCL5, MCP-1 and IL6 post treatment with each chemotherapy type. Doxorubicin monotherapy treated mice showed significantly longer overall survival compared to their carboplatin counterparts with further increases following addition of either STING agonist or PD-L1 ICB. However, despite the stronger ICD inducing ability of doxorubicin, overall survival of mice treated with carboplatin + STING agonist + PD-L1 ICB was the longest. Findings from our pre-clinical study provide novel insights for rationalized combinations of immune sensitizing agents such as STING pathway activators to improve response of HGSC patients to chemotherapy and ICB in the primary and recurrent settings.

## INTRODUCTION

High grade serous carcinoma of the ovary (HGSC) is a deadly disease with a five-year survival rate of just 45.6% that has seen little improvement over the past few decades^1, 2^. The poor survival rate of HGSC patients can be attributed to a variety of factors including diagnosis at later stages and high rates of resistance and recurrence^3^. Indeed, the majority of women diagnosed with HGSC present with advanced disease. At an advanced stage there are limited treatment options such as cytoreductive surgery followed by platinum and taxane-based combination chemotherapies. These treatments are largely ineffective as 70% of patients will relapse progressing to platinum-resistant HSGC. While other “hard-to-treat” tumor types have seen vast improvements with the integration of cancer immunotherapies, such as immune checkpoint blockade (ICB), HGSC has not seen the similar success^4^. Most ICB therapies have been shown to enhance the pre-existing immune landscape, specifically the presence of lymphocytes that express the target immune checkpoint for blocking and subsequent activation is required for effective treatment^5, 6^. For example, a pre-treatment tumour immune landscape with a high number of tumour infiltrating lymphocytes (TILs) is broadly defined as “hot/T-cell inflamed” and in most cases predictive of better prognosis and response to ICB when compared to their low TIL “cold/T cell non-inflamed” counterparts^7, 8^.

In our previous reports, we showed the pre-treatment immune transcriptome of tumours from platinum-resistant HGSC patients are intrinsically immunologically cold^9, 10^. We demonstrated that a non-inflamed pre-existing T helper type I tumor immune microenvironment (TIME), decreased expressions of type I interferon (IFN1) genes and STAT1 protein, low density of TILs associated with poor response to chemotherapy^10^. Strategies attempting to transform these immunologically cold tumors to hot, thus improving therapeutic response through IFN1 activation, have recently garnered tremendous interest across solid tumours^8, 11^.

One such example is using immune activating agents, including those that activate (cGAS)-Stimulator of Interferon Genes (STING) pathway (Figure 1A). Activation of STING pathway primarily occurs via cytosolic nucleic acid sensing that leads to increased production of IFN induced genes, specifically the TIL recruiting CXCR3 binding chemokines, CXCL9/10/11 and CCL5^11–13^. Supporting this hypothesis, we previously reported that response to platinum chemotherapy can be improved via incorporating STING agonist post carboplatin chemotherapy. In this pre-clinical study using the ID8-*Trp53^−/−^* syngeneic murine model of HGSC, we also showed the immune sensitizing potential of STING agonist to programmed cell death protein-1 (PD-1) ICB^12^. Addition of STING agonist to the treatment regimen significantly improved survival both when administered as a monotherapy and in combination with carboplatin and PD-1 ICB. Potentially attributed to the angiostatic function of CXCL10 chemokine, treatment with STING agonist also reduced ascites formation and overall tumor burden. Additionally, an enrichment of genes associated with antigen presentation, MHCII, IFN response, and increased expression of *Stat1* and *Cxcl10* leading to overall enhancement of IFN1 immune responses in the TIME, were observed in the tumours from mice treated with STING agonist^12^.

**Figure 1.**
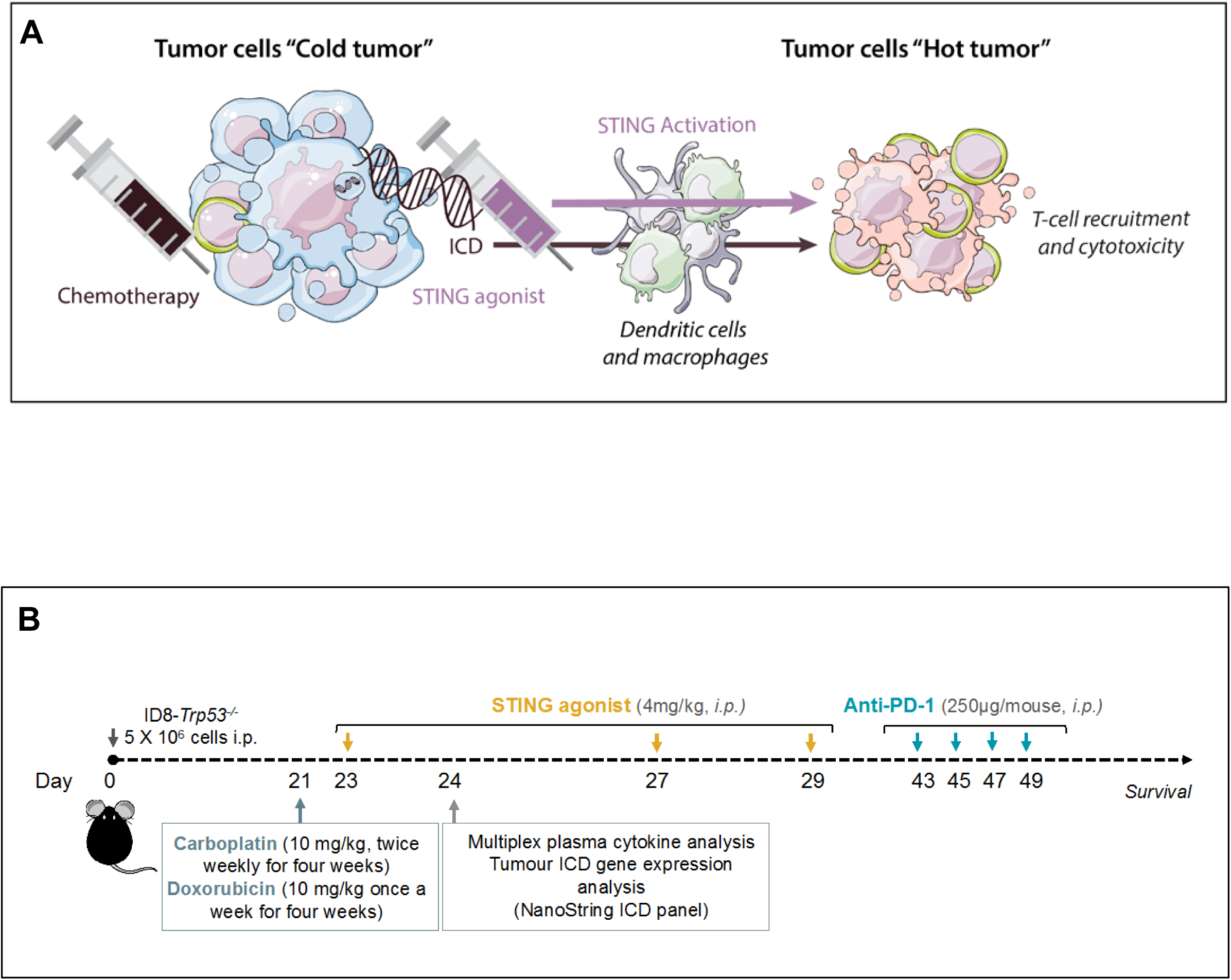
**A) Conceptual model for ICD potentiation via addition of STING agonist post chemotherapy treatment.** Response of an immunologically non-inflamed/cold tumour to treatment (chemotherapy/radiation and immune checkpoint blockade), could be enhanced by addition of STING agonist post chemotherapy. Chemotherapy specific immunogenic cell death (ICD) effect is key to response from STING activation and subsequent immune cell recruitment and antigen cross-presentation by myeloid cells post treatment. **B) Schematic showing the study design and treatment schedule in the ID8-*Trp53^−/−^* syngeneic murine model of HGSC.**

While these results provide a strong rationale for testing these combination treatment approaches for patients with platinum-sensitive tumors, as previously mentioned, many patients progress to develop platinum-resistant HGSC^1, 14^. Patients with recurrent HGSC are administered second line chemotherapies including doxorubicin, an anthracycline known to be a bonafide inducer of immunogenic cell death (ICD). ICD is an immune priming form of cell death which occurs following exposure to a subset of cytotoxic chemotherapies or radiotherapy^15–17^. Chemotherapy induced ICD can increase tumour antigen recognition and cross presentation by dendritic cells or macrophages to cytotoxic TILs in the sterile TIME. Following this logic, ICD inducing chemotherapies have been combined with ICB to further augment anti-tumor immunity^18^. In addition to their ICD inducing effects, anthracyclines, including doxorubicin, have also been reported to increase PD-L1 expression on tumor cells predicting a stronger rationale for combination with ICB anti-PD-L1 treatment^19, 20, 21^.

Based on these compelling findings, we sought to compare the efficacy of immune activating STING agonist when combined with a stronger ICD inducer – doxorubicin in the ID8-*Trp53^−/−^* model of HGSC. We hypothesized that the effects we previously reported with a combination of carboplatin and STING agonist, could be further potentiated with a stronger ICD inducer such as doxorubicin. Based on our previous finding that STING agonist treatment led to increased tumour and splenic myeloid derived suppressor cell specific PD-L1 expression, we further evaluated the impact of combination with PD-L1 ICB on overall survival. Findings from our study provide novel directions for the precise use of therapies activating STING pathway in combination with ICD inducing chemotherapy.

## METHODS

### Cell lines

The ID8-*Trp53^−/−^* mouse ovarian surface epithelial cells were kindly provided by Dr. Ian McNeish (Imperial College, London, UK)^21^. Mutations in *TP53* gene are present in >95% HGSC tumours^2^ and thus the recently modified ID8 cell line more closely recapitulates the human HGSC tumour progression. ID8-*Trp53^−/−^* cells were maintained in Dulbecco’s Modified Eagle’s Medium (Sigma Aldrich, Canada) supplemented with 4% fetal bovine serum, 100 µg/mL of penicillin/streptomycin and a solution containing 5 µg/mL of insulin, 2.5 µg/mL of transferrin and 2.5 ng/mL of sodium selenite.

### *In vivo* studies

All animal protocols were approved by the Queen’s University Animal Care Committee. 5-6 x 10^6^ ID8-*Trp53^−/−^* cells in 200 µl of PBS were transplanted via intra-peritoneal (IP) injections in eight to ten-week old female C57BL/6 mice (Charles River Laboratories International Inc). Approximately 4 weeks post tumour cell implantation, mice were randomized and treated with 1) Carboplatin, 2) Doxorubicin, 3) Carboplatin + STING agonist, 4) Doxorubicin + STING agonist 5) Carboplatin + STING agonist + anti-PD-L1 or 6) Doxorubicin + STING agonist + anti-PD-L1 via IP administration, at the indicated doses and time points (Figure 1B). The anti-mouse PD-L1 antibody (clone RMP1-14; BioXcell) was administered two weeks following the last STING agonist injection at a dose of 200 µg per animal at two-day intervals for a total of four injections via IP route.

### Plasma cytokine profiling

To determine the effect of chemotherapy type and combination with STING agonist, on the systemic cytokine profiles, plasma samples collected at 24 h time point following first STING agonist treatment post either carboplatin or doxorubicin treatment, were subjected to multiplexed cytokine profiling using the MD31, 31-plex cytokine/chemokine array (Eotaxin, G-CSF, GM-CSF, IFN gamma, IL-1alpha, IL-1beta, IL-2, IL-3, IL-4, IL-5, IL-6, IL-7, IL-9, IL-10, IL-12 (p40), IL-12 (p70), IL-13, IL-15, IL-17A, IP-10, CXCL1, LIF, LIX, MCP-1, M-CSF, MIG, MIP-1alpha, MIP-1beta, MIP-2, RANTES, TNF alpha, VEGF) at Eve Technologies Corporation (Calgary, AB, Canada). All samples were analysed in biological triplicates. The standard curve regression was used to calculate the concentration of each target cytokine. Differences between levels of cytokines were analysed using GraphPad Prism (7.02).

### Tumour ICD gene expression profiling using a custom NanoString panel

To determine the ICD effect induced by carboplatin and doxorubicin chemotherapy and subsequent effects post addition of STING agonist, total RNA from tumours collected 24 h post first STING agonist treatment and the chemotherapy only controls, A were subjected to NanoString based gene expression profiling using a custom ICD gene panel (Table 1). Briefly, total RNA from fresh frozen tumour tissues was isolated using the total RNA Purification Kit (Norgen Biotek Corporation) as per the manufacturer’s instructions. RNA concentration and purity were estimated on a NanoDrop ND-100 spectrophotometer (NanoDrop Technologies, Wilmington, DE, USA). 150 ng of total RNA from each tumour sample was subjected to digital multiplexed profiling, using a custom ICD gene panel (60 ICD related genes with 5 housekeeping controls, NanoString Technologies Inc.) as per our previously established protocols. Normalization of raw data was performed using the nSolver software 3.0 (NanoString Technologies, Seattle, WA). The raw NanoString counts were initially subjected to normalization for all target RNAs in all samples based on built-in positive controls. This step accounts for inter-sample and experimental variation such as hybridization efficiency and post-hybridization processing. The geometric mean of each control was then calculated to indicate the overall assay efficiency. The housekeeping genes were used for mRNA content normalization. Differentially expressed genes between the tumours from different treatment groups were derived using GraphPad Prism software. A p-value <0.05 was considered statistically significant.

**Table 1.**
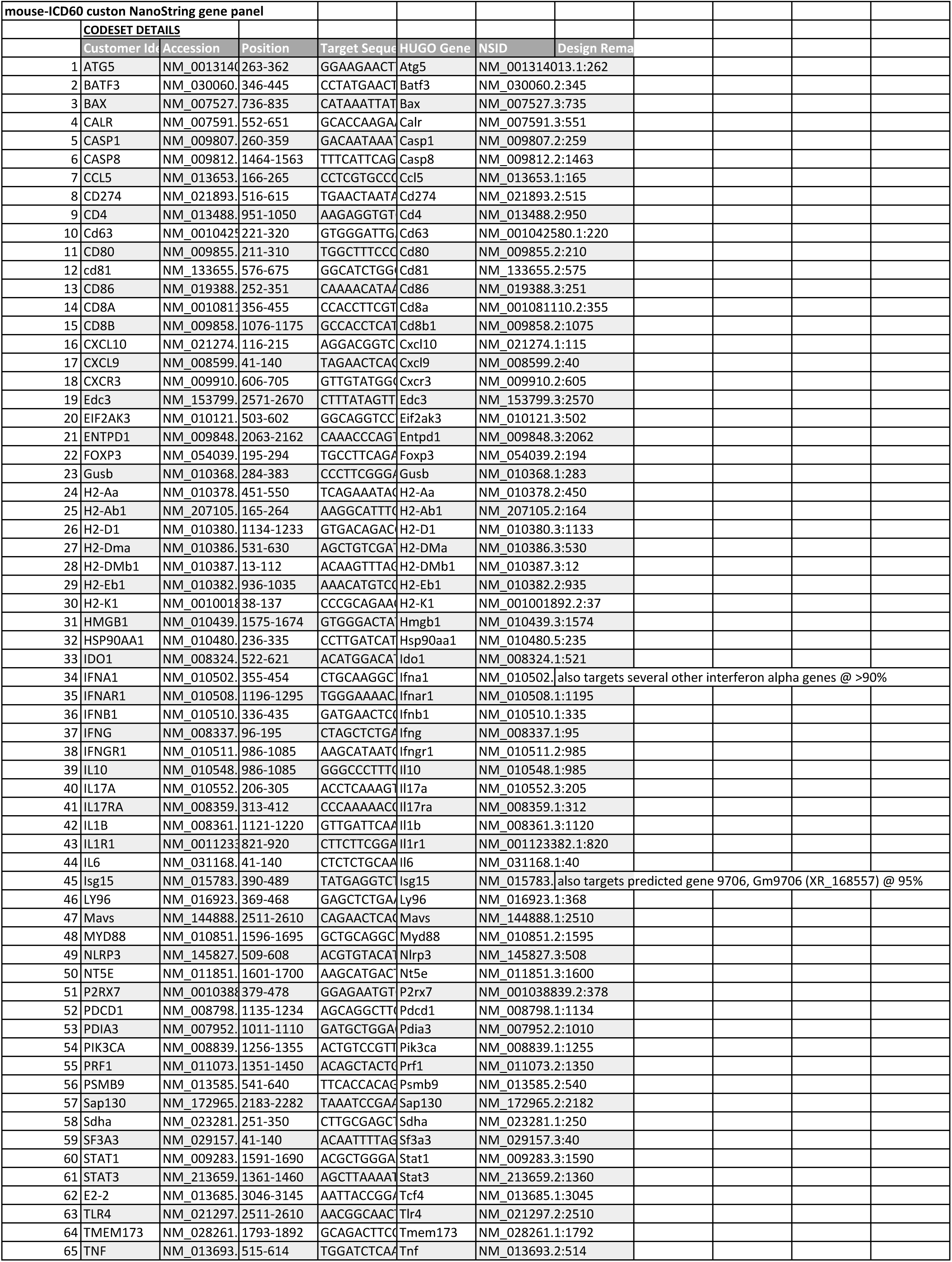
Custom ICD gene panel for NanoString platform based gene expression profiling.

## RESULTS

### Doxorubicin induces a higher and distinct expression of ICD genes compared to carboplatin in ID8-*Trp53^−/−^* tumours

The expression of 60 ICD associated genes was measured in RNA isolated from tumours of mice treated with carboplatin or doxorubicin monotherapy (Figure 2A and 2B). Genes specifically associated with IFN1 pathways, including, *Ifna1, Ifnb1, Psmb9 and Cxcl10*, showed significantly (p<0.05) higher expression in doxorubicin treated mice compared to those treated with carboplatin (Figure 2B). Interestingly, *Ccl5* expression was higher in carboplatin treated tumours compared to those treated with doxorubicin.

**Figure 2A.**
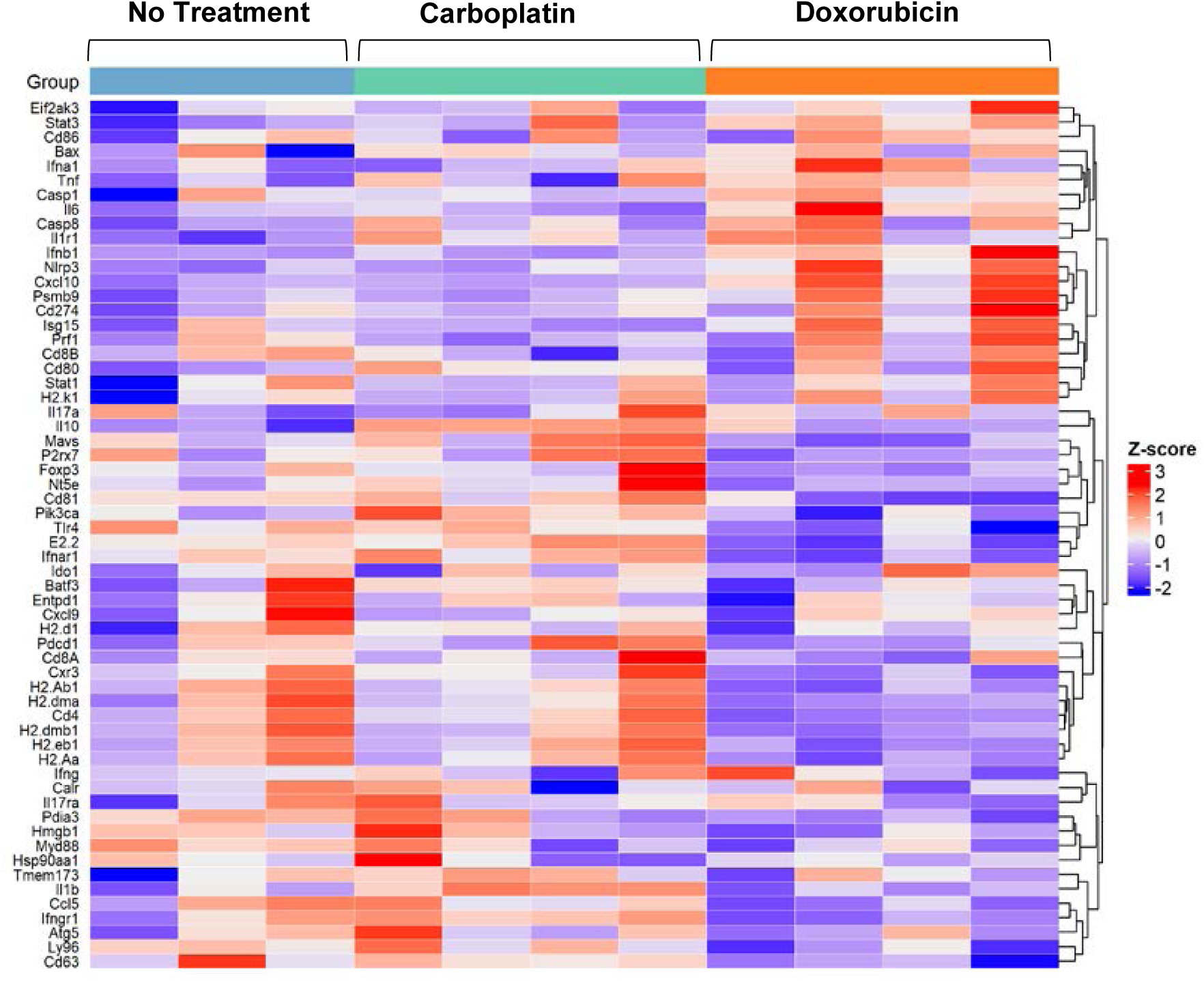
Doxorubicin activates higher immune responses within the tumour immune microenvironment compared to carboplatin. Heat map showing expression pattern of 60 ICD associated genes in tumours from mice treated with carboplatin compared to doxorubicin. Scale function was used to center the expression scores and ComplexHeatmap package was used to build the heatmaps in R Bioconductor statistical environment.

**Figure 2B.**
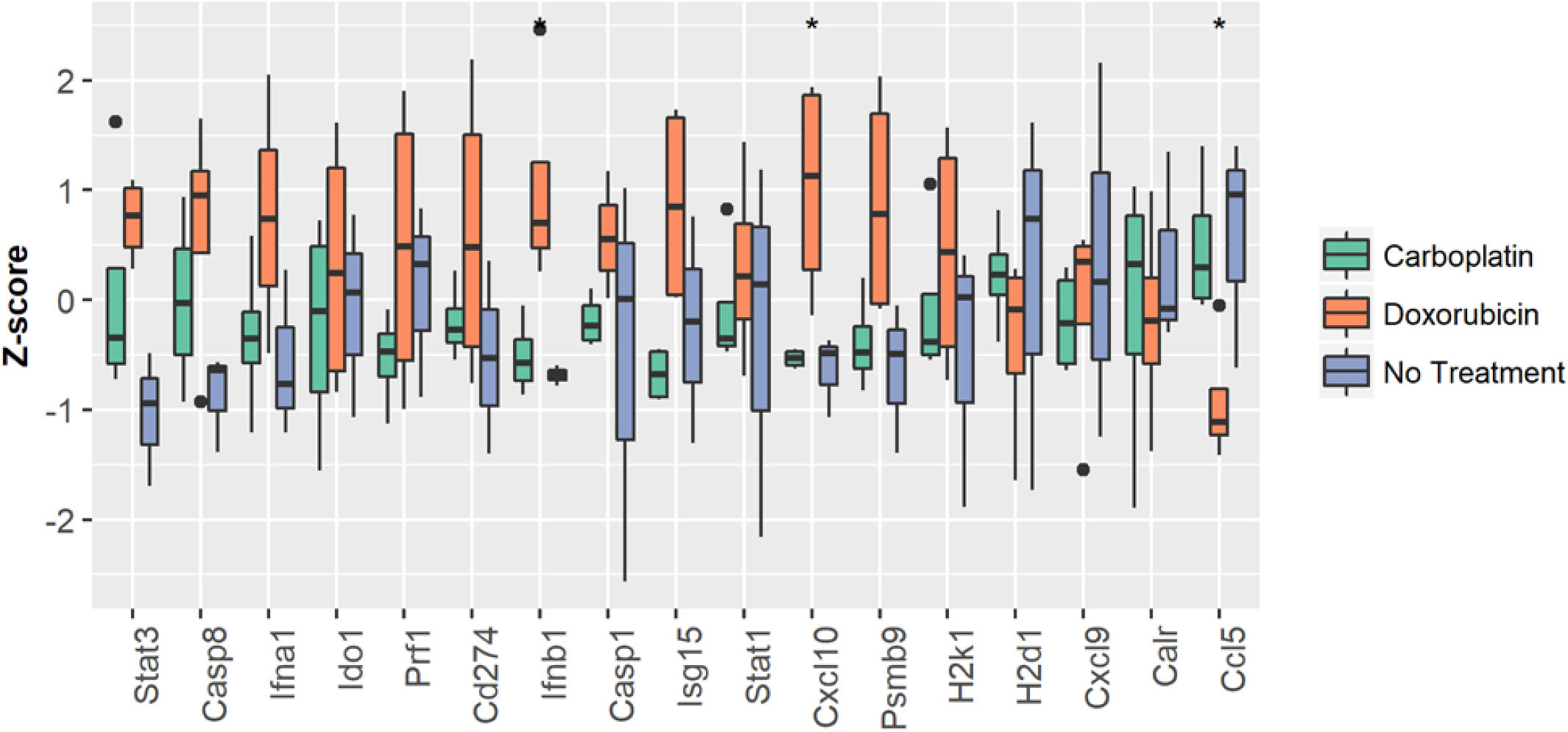
Doxorubicin induces differential ICD gene expression compared to carboplatin. A 60 gene custom ICD NanoString panel was applied to measure the expression of genes associated with ICD pathways. Kruskal Wallis test was applied to compare the median between the three groups. Data analysis was performed using R Bioconductor. p-value<0.05 (*) was considered statistically significant.

### STING activation alters expression of tumour ICD associated genes in a chemotherapy specific manner

Addition of STING agonist post chemotherapy showed significant differences in the magnitude of ICD gene expression in tumours (Figure 3A). In general, doxorubicin + STING agonist combination showed significantly (p<0.05) higher expression of *Stat3, Casp8, Ifna1, Ido1, Prf1, CD274, Ifnb1, Casp1, Isg15, Stat1, Cxcl10, Psmb9, H2k1* and *H2d1*, compared to carboplatin chemotherapy (Figure 3B). Interestingly, however, the expression of *Cxcl9, Calr* and *Ccl5* was significantly higher in carboplatin + STING agonist treated tumours compared to those treated with doxorubicin (Figure 3B) indicative of their possible differential expression in cancer cells compared to immune cells.

**Figure 3A.**
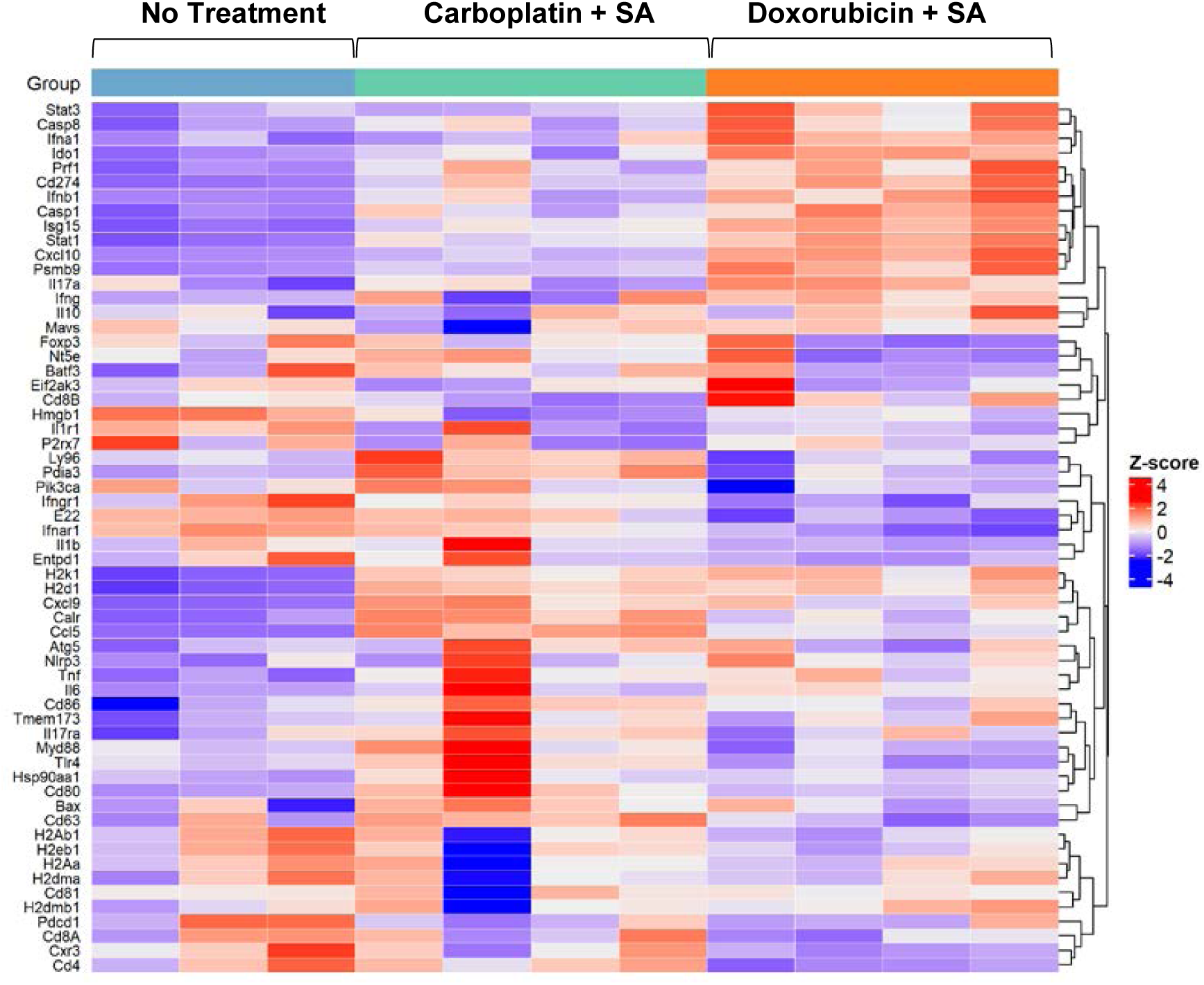
STING agonist potentiates doxorubicin induced ICD. Heat map showing differential expression pattern of 60 ICD associated genes in tumours from mice treated with carboplatin + STING agonist (SA) compared to doxorubicin + SA. Scale function was used to center the expression scores and ComplexHeatmap package was used to build the heatmaps in R.

**Figure 3B.**
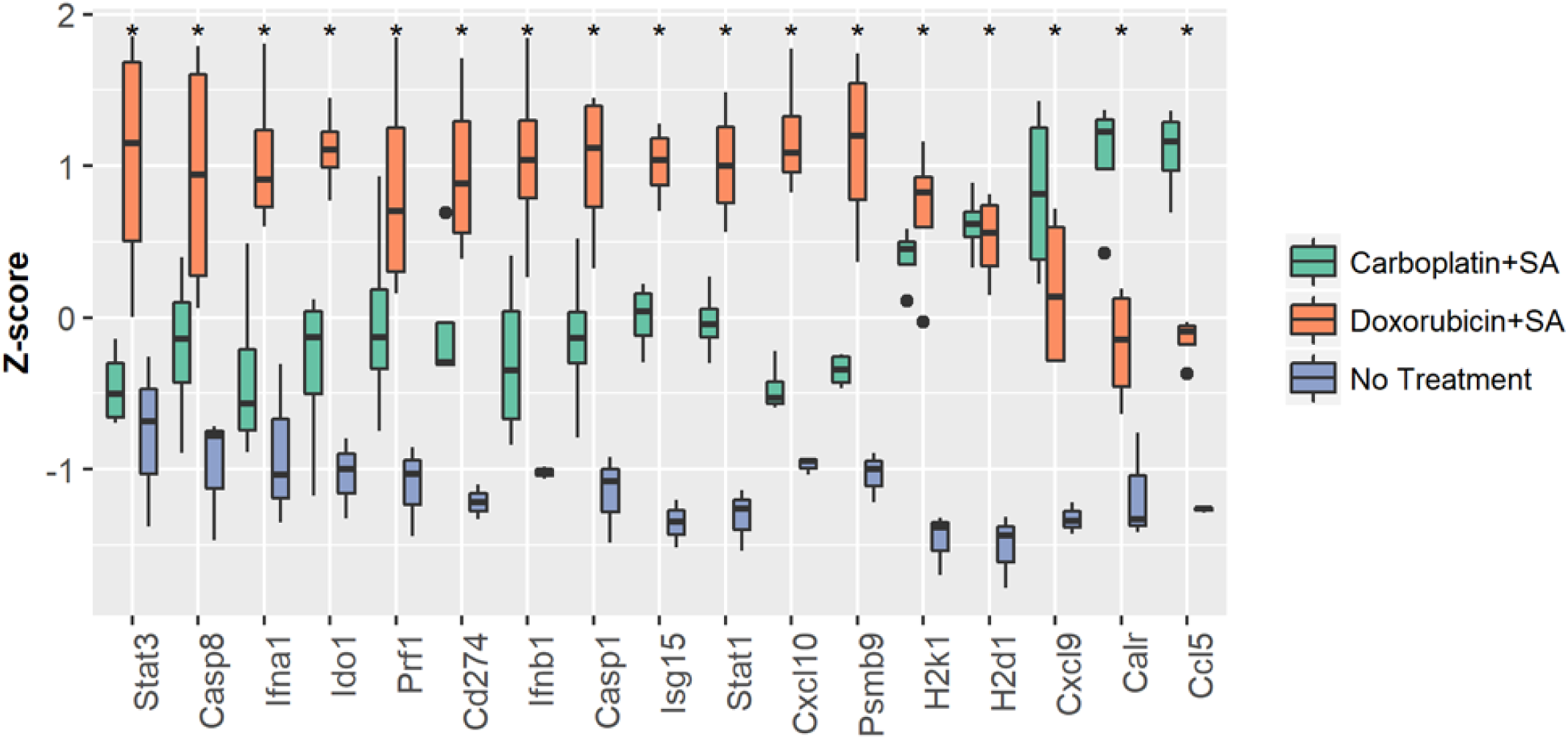
STING agonist affects the expression of ICD genes in chemotherapy specific manner. Tumours from doxorubicin + STING agonist (SA) treated mice and carboplatin + SA treated mice were subjected to ICD gene expression profiling using a custom NanoString panel. Kruskal Wallis test was applied to compare the median between the three groups. Data analysis was performed using R Bioconductor. p-value<0.05 (*) was considered statistically significant.

### STING agonist amplifies doxorubicin mediated cytokine production

To determine the chemotherapy associated differences in plasma cytokine levels, we conducted multiplex cytokine analysis of plasma collected at 24 h post first treatment with carboplatin, doxorubicin or untreated controls. Doxorubicin only treated mice showed elevated plasma levels if CXCL10, MCP-1, MIP-1B compared to those treated with carboplatin, however this difference was not statistically significant (Figure 4). Notably, the levels of CCL5 and IL-6 were significantly higher in doxorubicin treated mice compared to those treated with carboplatin (Figure 4).

**Figure 4.**
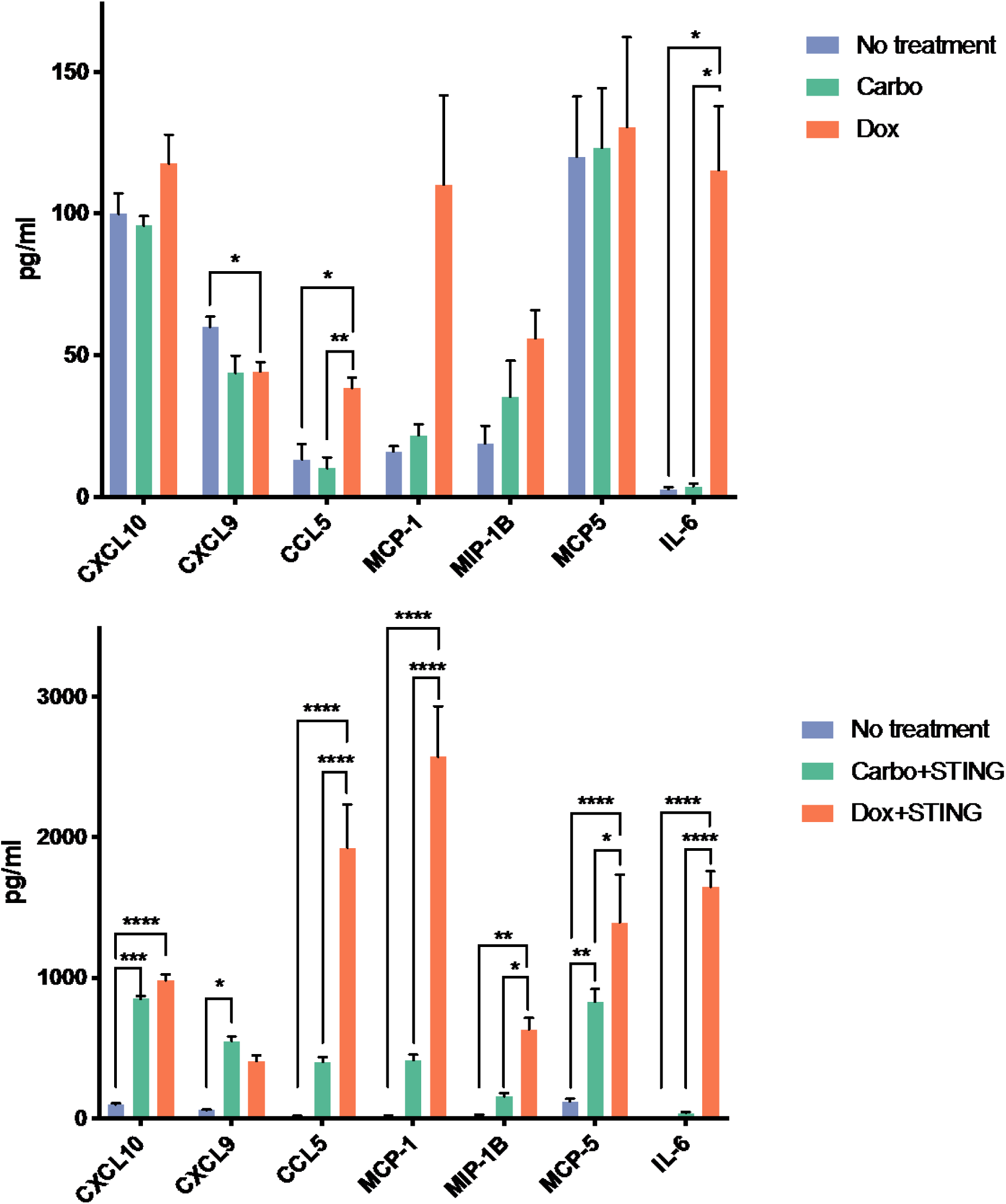
Distinct plasma cytokine profile observed in mice treated with doxorubicin or carboplatin chemotherapy is further amplified with the addition of STING agonist. ID8-*Trp53*^−/−^ tumor bearing mice were treated with **(A)** control, carboplatin or doxorubicin and **(B)** carboplatin + STING agonist or doxorubicin + STING agonist. Two-way ANOVA with Tukey’s post hoc test was performed using GraphPad prism (mean +/− SEM, \**P*<0.05, \*\**P*<0.01, \*\*\**P*<0.001, \*\*\*\**P*<0.0001)

The addition of STING agonist further elevated CXCL10 levels in the plasma of carboplatin and doxorubicin treated mice, however the difference between the two chemotherapy types was not significant (Figure 4). Interestingly, addition of STING agonist led to significantly increased levels of CXCL9 in carboplatin treated mice only (Figure 4). STING agonist treatment also further amplified CCL5 levels (p < 0.0001) in doxorubicin treated mice compared to both carboplatin + STING agonist and vehicle control groups (Figure 4). Similar response patterns were observed in levels of MCP-1, MIP-1B, MCP-5 and IL-6.

### Addition of STING agonist post doxorubicin chemotherapy does not add a survival advantage compared to carboplatin

To evaluate the differential impact on overall survival, doxorubicin and carboplatin were used as single agents or in combination with a) STING agonist, b) anti-PD-L1 ICB or c) STING agonist and anti-PD-L1, in the ID8-*Trp53^−/−^* syngeneic mouse model. The rationale for addition of PD-L1 ICB was based on post treatment tumour gene expression profiling that showed increased levels of *Cd274* (gene encoding PD-L1) following addition of STING agonist.

Comparison of chemotherapy types as single agents revealed that doxorubicin treated mice had significantly longer overall survival (average of 96.5 days) than carboplatin treated mice (average of 77 days; Figure 5). In line with our previously reported findings, addition of STING agonist significantly increased the survival of carboplatin treated mice by an average of 13 days (Figure 5). Although, the addition of anti-PD-L1 to carboplatin did not show any significant increase in overall survival, treatment with anti-PD-L1 following treatment with STING agonist and carboplatin chemotherapy significantly extended the median overall survival to 101 days (Figure 5B).

**Figure 5.**
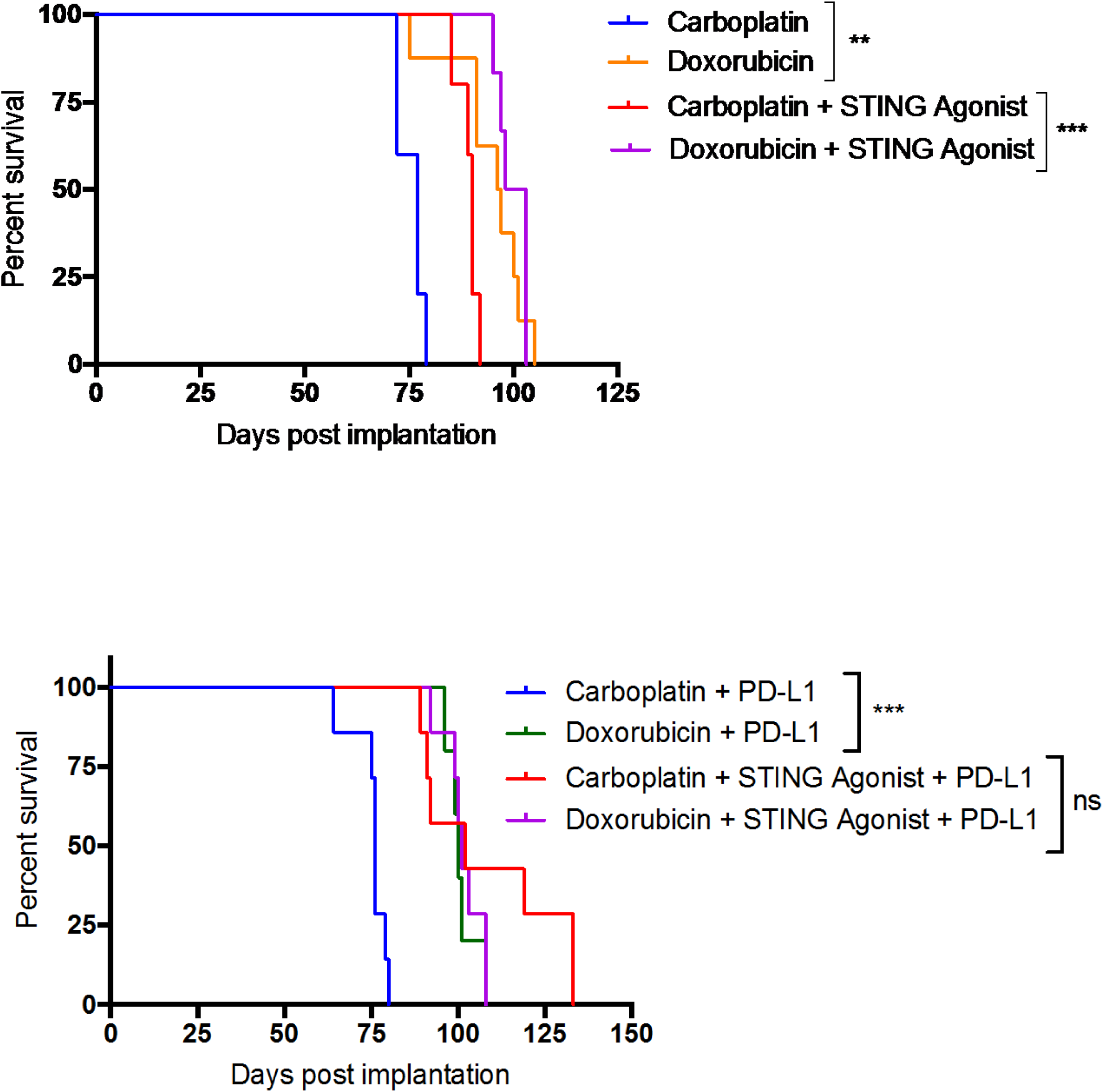
Effect of STING agonist on overall survival and immune sensitization to PD-L1 ICB in combination with carboplatin or doxorubicin chemotherapy. Kaplan Meier survival analysis was performed using Graphpad prism software (version 7.0). Log-rank Mantel cox test was applied to determine statistically significant differences. p-value<0.05 (*) was considered statistically significant; ** *P*<0.01, \*\*\**P*<0.001)

Surprisingly, upon addition of STING agonist or anti-PD-L1, we did not observe significantly increased survival advantage in the doxorubicin treated mice, with a modest increase in survival of 4 and 3.5 days respectively. Overall survival of doxorubicin + STING + anti-PD-L1 combination was, however, significantly increased to an average of 103 days (Figures 5A and B).

## DISCUSSION

Activation of the cGAS-STING pathway, via direct (STING agonist, oncolytic virus) and indirect (radiation, PARP inhibitors) approaches, is an emerging immune adjuvant treatment approach^8, 11^. With several promising pre-clinical findings across an array of cancer models, recent reports including ours have confirmed the immune sensitizing effect of STING pathway activation thus improving response to conventional chemotherapy and novel ICB^12^. Importantly, our previous report showed that direct activation of STING pathway can enhance response to carboplatin chemotherapy and sensitize tumours to PD-1 ICB.

Patients with HGSC have minimally benefited from the newer ICB therapies indicating the lack of understanding in how the effect of ICB is dependent on the pre-existing TIME^22^. It is well established that HGSC tumours exhibit high genomic instability and thus are thought to be immunogenic, contributing to immunologically variant states that can be identified at the time of diagnosis or primary debulking surgery^23^. In line with these characteristics, we and others have previously shown that pre-existing immunologically divergent states also associate with chemotherapy response and overall survival, indicating the significance of IFN induced chemokines and associated ICD in mediating treatment response^10, 24^. It is important to note that a growing body of evidence suggests a co-existence of inflamed and non-inflamed states across multiple regions in one given tumour^25^. However, irrespective of their classification, the density, localization and activation state of immune cells in the TIME could greatly impact variability in therapeutic response to immune based therapies, in addition to the type of chemotherapy^26, 27^.

The level of ICD response elicited by chemotherapies is one example of a mechanism dependent on the pre-existing TIME. Specifically, ICD events lead to a release in danger associated molecular pattern (DAMP) in a spatiotemporal manner that can have a profound impact on the consequent activation of both innate and adaptive immune response. Therefore, a comprehensive understanding of treatment induced ICD events is critical for the design of rationalistic ICB combinations^28, 29^. While radiation is the most potent ICD inducing therapy, chemotherapeutic agents such as anthracyclines, platinum, taxanes and other agents promote variable degrees of ICD events that ultimately alter tumour immunogenicity^30, 31^. Along this notion, it can be speculated that inflamed tumours with high pre-existing activated TILs and IFN gene expression will produce a higher magnitude of immune-mediated responses compared to non-inflamed tumours given the proximity of intra-tumourally located TILs. Indeed, ongoing ICB combination trials are exploring radiation induced ICD led immune sensitization in solid tumours. In the context of HGSC, PARP inhibitor induced STING pathway activation and combination with ICB is under several clinical trials^32^.

In HGSC, platinum-taxane based chemotherapy is widely used in the frontline setting whereas liposomal doxorubicin is practiced in post recurrence treatment in the second line setting^33^. Carboplatin and doxorubicin, elicit their cancer cell killing effects via distinct modes of action. For example, carboplatin functions by eliciting DNA damage to block replicative machinery and ultimately causes the cell to undergo apoptosis while doxorubicin intercalates with DNA to inhibit topoisomerase II function and produces a high level of reactive oxygen species leading to membrane damage^34, 35^. While both are known to induce ICD, they achieve cell death through differing molecular mechanisms resulting in varying levels of ICD. Several previous reports have exploited these distinct capacities of doxorubicin with regard to cellular IFNI responses^17^. Most recently Wilkinson et al., show this effect as a result of differential activation of cGAS-STING pathway in a chemotherapy specific manner (https://www.biorxiv.org/content/10.1101/764662v2). This study demonstrates the significant increase in levels of CXCL10/CCL5 as result of release of micronuclei following treatment with doxorubicin.

Towards their rationalized clinical translation in HGSC and differences in chemotherapy specific ICD inducing ability, in the current study, we evaluated the effect of synergistic STING pathway activation in the context of carboplatin and doxorubicin chemotherapy in the ID8-*Trp53*^−/−^ syngeneic murine model of ovarian cancer. In concordance with the findings by Wilkinson et al. 2019 and others with regard to doxorubicin associated cGAS-STING activation, we observed increases in plasma CXCL10/CCL5 levels post doxorubicin treatment compared to carboplatin. Surprisingly, the expression of *Ccl5* gene was significantly higher in carboplatin and carboplatin + STING agonist treated tumours compared to those treated with doxorubicin and doxorubicin +STING agonist. This finding is suggestive of potential differences in cancer cell and immune cell specific STING pathway activation and warrants further investigation.

When used as a single agent, doxorubicin treated mice had a significant increase in survival compared to those treated with carboplatin. Similar to our previous findings, the significant increase in survival following addition of STING agonist to either carboplatin or doxorubicin treated tumours strongly suggests that immunomodulation via STING pathway activation post chemotherapy could be a promising approach to improve response to carboplatin chemotherapy. Surprisingly, survival was not further prolonged following the addition of STING agonist in the doxorubicin treated mice, and therefore we observed for the first time that response to doxorubicin treatment may not achieve the level of improvement as seen with carboplatin from the addition of STING agonist. Our leading explanation for this finding is the differential ICD response produced by doxorubicin and carboplatin^36^. Doxorubicin itself is known to induce IFN1 response via STING pathway activation and downstream chemokine induction and therefore the addition of STING agonist may not significantly increase immune activation in the doxorubicin treated TIME^37^.

Tumour immune transcriptomic profiles 24 hours post chemotherapy and STING agonist treatment showed significant increase in expression of *Cd274* (gene encoding PD-L1). Furthermore, in our previous report we observed increased levels of PD-L1 in splenic myeloid derived suppressor cells post addition of STING agonist to carboplatin. We thus added anti-PD-L1 to the treatment regimen. Interestingly, the addition of anti-PD-L1 ICB to the doxorubicin + STING agonist treated group did not add further survival benefit. This was indeed an unexpected finding given the large impact it had on survival following carboplatin treatment. However, doxorubicin is known to impact PD-L1 expression, such as decreased surface expression and increased nuclear expression on breast cancer cells^38^. This altered expression could potentially have impacted the response to anti-PD-L1 therapy in this study. Another possible reason could be the increased IL-6 level post doxorubicin treatment, which was amplified by a magnitude of >15 fold upon addition of STING agonist. As previously reported, the significant increase in IL6 might have contributed to lack of survival benefit in these mice due to its immunosuppressive effects on CD4+ T cells and increasing cancer cell PD-L1 expression^39^. IL-6 promotes the survival of cancer cells and is usually associated with poor prognosis across cancer types. Importantly, in the context of immunotherapies, high IL-6 level is the key indicator of cytokine release syndrome that is usually associated with immune related adverse events^40, 41^. Moreover, IL-6 is well known to be associated with chemotherapy resistance in HGSC, as previously reported by us and others^10^. As recently shown in melanoma models, blockade of IL-6 following STING agonist treatment post doxorubicin treatment might prolong survival via augmenting T helper type I response^39^. However, a significant increase in survival was observed when doxorubicin, STING agonist and anti-PD-L1 were used in combination. Blockade of IL6 in these mice post addition of STING agonist may potentially lead to an increased survival benefit.

With growing awareness that the TIME is both impacted by and impacts the efficacy of cancer therapies, it’s imperative that combination immunotherapy strategies are rationally designed. This study is the first to directly compare the combination of STING agonists with differential chemotherapies within the same model and importantly identifies that chemotherapy combinations with STING can mimic the TIME effects of a strong ICD inducer for precise immune sensitization for PD-L1 ICB. This key finding suggests that in the development of combination immunotherapeutic strategies to avoid high-dose toxicity associated with chemotherapies, other agents inducing immune-stimulating pathways can be co-administered to prevent toxicities while eliciting similar immune stimulating effects.

Our study significantly advances the field of STING pathway activation based immune sensitization of tumours, however, there are some limitations to our study design. Since our question was primarily to evaluate *in vivo* differences in the synergistic effect of STING activation with different chemotherapy types and PD-L1 ICB, we did not perform the gold standard ICD induction assay in cancer cells prior to implantation in mice, as suggested by the consensus ICD guidelines proposed by Kepp et al.,^42^. Indeed, results from our study warrant future mechanistic studies to determine the cancer cell vs immune cell effects of STING activation following ICD inducing therapies. In conclusion, our novel findings, form the basis for rationalized combinations of STING pathway activation to improve chemotherapy response and sensitize HGSC to PD-L1 ICB.

## Acknowledgements

MK conceptualized and designed the study. SN, NS and MK wrote the manuscript. TV reviewed and conducted the statistical analysis of NanoString data. SN, NS and EL conducted the experiments. AA performed statistical analysis of cytokine data. All co-authors reviewed the manuscript. This study was funded by grants from the Canadian Institutes of Health Research and Ontario Ministry of Research Innovation and Science; Early Researcher Award to MK.

## References

1. Wilson, M. K. et al. Fifth ovarian cancer consensus conference of the gynecologic cancer intergroup: Recurrent disease. Ann. Oncol. 28, 727–732 (2017).

2. Clifford, C. et al. Multi-omics in high-grade serous ovarian cancer: Biomarkers from genome to the immunome. Am. J. Reprod. Immunol. (2018). doi:10.1111/aji.12975

3. Bosquet, J. G., Marchion, D. C., Chon, H., Lancaster, J. M. & Chanock, S. Analysis of chemotherapeutic response in ovarian cancers using publicly available high-throughput data. Cancer Res. 74, 3902–3912 (2014).

4. Odunsi, K. Immunotherapy in ovarian cancer Symposium article. 28, 1–7 (2017).

5. Dallos, M. C. & Drake, C. G. Blocking PD-1/PD-L1 in Genitourinary Malignancies. Cancer J. (United States) 24, 20–30 (2018).

6. Pitt, J. M. et al. Resistance Mechanisms to Immune-Checkpoint Blockade in Cancer: Tumor-Intrinsic and -Extrinsic Factors. Immunity 44, 1255–1269 (2016).

7. Danaher, P. et al. Pan-cancer adaptive immune resistance as defined by the Tumor Inflammation Signature (TIS): Results from The Cancer Genome Atlas (TCGA). J. Immunother. Cancer 6, 1–17 (2018).

8. Galon, J. & Bruni, D. Approaches to treat immune hot, altered and cold tumours with combination immunotherapies. Nat. Rev. Drug Discov. 18, 197–218 (2019).

9. Koti, M. et al. A distinct pre-existing inflammatory tumour microenvironment is associated with chemotherapy resistance in high-grade serous epithelial ovarian cancer. Br. J. Cancer 112, 1215–1222 (2015).

10. Au, K. K. et al. STAT1-associated intratumoural T _H_ 1 immunity predicts chemotherapy resistance in high-grade serous ovarian cancer. J. Pathol. Clin. Res. (2016). doi:10.1002/cjp2.55

11. Flood, B. A., Higgs, E. F., Li, S., Luke, J. J. & Gajewski, T. F. STING pathway agonism as a cancer therapeutic. Immunol. Rev. 290, 24–38 (2019).

12. Ghaffari, A. et al. STING agonist therapy in combination with PD-1 immune checkpoint blockade enhances response to carboplatin chemotherapy in high-grade serous ovarian cancer. Br. J. Cancer (2018). doi:10.1038/s41416-018-0188-5

13. Ramanjulu, J. M. et al. Design of amidobenzimidazole STING receptor agonists with systemic activity. Nature 564, 439–443 (2018).

14. Jayson, G. C., Kohn, E. C., Kitchener, H. C. & Ledermann, J. a. Ovarian cancer. Lancet 384, 1376–1388 (2014).

15. Kroemer, G., Galluzzi, L., Kepp, O. & Zitvogel, L. Immunogenic cell death in cancer therapy. Annu. Rev. Immunol. 31, 51–72 (2013).

16. Fridman, W. H., Zitvogel, L., Sautès-Fridman, C. & Kroemer, G. The immune contexture in cancer prognosis and treatment. Nat. Rev. Clin. Oncol. 14, 717–734 (2017).

17. Sistigu, A. et al. Cancer cell-autonomous contribution of type I interferon signaling to the efficacy of chemotherapy. Nat. Med. 20, 1301–1309 (2014).

18. Obeid, M. et al. Calreticulin exposure dictates the immunogenicity of cancer cell death. Nat. Med. 13, 54–61 (2007).

19. Yoon, H. K. et al. Effect of anthracycline and taxane on the expression of programmed cell death ligand-1 and galectin-9 in triple-negative breast cancer. Pathol. Res. Pract. 214, 1626–1631 (2018).

20. Wang, J. et al. Checkpoint Blockade in Combination With Doxorubicin Augments Tumor Cell Apoptosis in Osteosarcoma. J. Immunother. 42, 321–330 (2019).

21. Walton, J. et al. CRISPR / Cas9-Mediated Trp53 and Brca2 Knockout to Generate Improved Murine Models of Ovarian High-Grade Serous Carcinoma. 2000, 6118–6129 (2016).

22. Gibney, G. T., Weiner, L. M. & Atkins, M. B. Predictive biomarkers for checkpoint inhibitor-based immunotherapy. Lancet Oncol. 17, e542–e551 (2016).

23. Bowtell, D. D. et al. Rethinking ovarian cancer II: reducing mortality from high-grade serous ovarian cancer. Nat. Rev. Cancer 15, 668–679 (2015).

24. Au, K. K. et al. Gynecologic Oncology CXCL10 alters the tumour immune microenvironment and disease progression in a syngeneic murine model of high-grade serous ovarian cancer. Gynecol. Oncol. 4–13 (2017). doi:10.1016/j.ygyno.2017.03.007

25. Zhang, A. W. et al. Interfaces of Malignant and Immunologic Clonal Dynamics in Ovarian Cancer. Cell 173, 1755–1769.e22 (2018).

26. Cristescu, R. et al. Pan-tumor genomic biomarkers for PD-1 checkpoint blockade-based immunotherapy. Science (80-.). 362, (2018).

27. Mariathasan, S. et al. TGFβ attenuates tumour response to PD-L1 blockade by contributing to exclusion of T cells. Nature 554, 544–548 (2018).

28. Lo, C. S. et al. Neoadjuvant chemotherapy of ovarian cancer results in three patterns of tumor-infiltrating lymphocyte response with distinct implications for immunotherapy. Am. Assoc. Cancer Res. clincanres.1433.2016 (2016). doi:10.1158/1078-0432.ccr-16-1433

29. Kroemer, G., Galluzzi, L., Kepp, O. & Zitvogel, L. Immunogenic Cell Death in Cancer Therapy ICD: immunogenic cell death. Annu. Rev. Immunol 31, 51–72 (2013).

30. Solanki, A. A. et al. Combining Immunotherapy with Radiotherapy for the Treatment of Genitourinary Malignancies. Eur. Urol. Oncol. 2, 79–87 (2019).

31. Rodríguez-Ruiz, M. E., Vanpouille-Box, C., Melero, I., Formenti, S. C. & Demaria, S. Immunological Mechanisms Responsible for Radiation-Induced Abscopal Effect. Trends Immunol. 39, 644–655 (2018).

32. Ding, L. et al. PARP Inhibition Elicits STING-Dependent Antitumor Immunity in Brca1-Deficient Ovarian Cancer. Cell Rep. 25, 2972–2980.e5 (2018).

33. Blake, E. A. et al. Efficacy of pegylated liposomal doxorubicin maintenance therapy in platinum-sensitive recurrent epithelial ovarian cancer: a retrospective study. Arch. Gynecol. Obstet. 299, 1641–1649 (2019).

34. Aletras, V., Hadjiliadis, D. & Hadjiliadis, N. Elements of the mechanism of action of the antitumor drug cis-platin or cis-DDP and its second generation derivatives. Ep. Klin. Farmakol. kai Farmakokinet. 13, 153–180 (1995).

35. Fucikova, J. et al. Human tumor cells killed by anthracyclines induce a tumor-specific immune response. Cancer Res. 71, 4821–4833 (2011).

36. Kepp, O. & Kroemer, G. Combinatorial strategies for the induction of immunogenic cell death. 6, 1–11 (2015).

37. Ii, T. & Induce, I. crossm. 8, (2017).

38. Ghebeh, H. et al. Doxorubicin downregulates cell surface B7-H1 expression and upregulates its nuclear expression in breast cancer cells: Role of B7-H1 as an anti-apoptotic molecule. Breast Cancer Res. 12, (2010).

39. Tsukamoto, H. et al. Combined blockade of IL6 and PD-1/PD-L1 signaling abrogates mutual regulation of their immunosuppressive effects in the tumor microenvironment. Cancer Res. 78, 5011–5022 (2018).

40. Rooney, C. & Sauer, T. Modeling cytokine release syndrome. Nat. Med. 24, 705–706 (2018).

41. Teachey, D. T. et al. Identification of predictive biomarkers for cytokine release syndrome after chimeric antigen receptor T-cell therapy for acute lymphoblastic leukemia. Cancer Discov. 6, 664–679 (2016).

42. Kepp, O. et al. Consensus guidelines for the detection of immunogenic cell death. Oncoimmunology 3, 1–19 (2014).

